# Parabens Impair Mouse Antral Follicle Growth and Alter Ovarian Hormone and Gene Expression Similar to Phthalates

**DOI:** 10.64898/2026.10.01.756089

**Authors:** Courtney Potts, Arena Chowdhury, Raulle Reynolds, Joao Agudo, Stephanie Martinez, Hanin Alahmadi, Genoa R. Warner

## Abstract

Benzyl paraben (BzP) and butyl paraben (BuP) are used as preservatives and antimicrobial agents in cosmetics, pharmaceuticals, and foods. BzP and BuP have been detected in urine, placental tissue, and ovarian follicular fluid and are understudied with regards to their endocrine activity, raising concerns on their impact to the reproductive system. We tested the hypothesis that BzP and BuP are toxic to the ovary and behave similarly to other structurally similar endocrine disrupting chemicals, such as phthalates and terephthalates, by altering hormone levels and impacting cell growth and gene expression in vitro. Untreated mouse antral follicles were exposed to vehicle, 0.1, 1, 10, or 100 µg/mL BzP, BuP, or the BuP analog monobutyl terephthalate (MBuTP) for 96 hours. The highest dose of BzP and BuP were cytotoxic to follicles, suggesting that medium chain length hydroxybenzoates are more cytotoxic to the ovary than corresponding terephthalic acids. BzP significantly increased levels of estradiol and testosterone compared to control. BuP decreased follicle growth and disrupted expression of cell cycle regulators, similar to phthalates. The peroxisome proliferator-activated receptor γ (PPARγ) transcriptionally controlled gene *Fabp4* was significantly upregulated by both BzP and BuP, whereas MBuTP upregulated all four PPAR target genes and decreased follicle growth. Overall, BzP, BuP, and MBuTP disrupted hormone levels, follicle growth, and cell cycle progression, suggesting similarities in toxicity to phthalates and potential endocrine disruption. Further study should be done on the reproductive toxicity and endocrine disrupting effects of chemicals similar in structure to phthalates to reduce exposure to endocrine disruptors and inform design of safer replacements.

## 1. Introduction

Endocrine-disrupting chemicals (EDCs) are prevalent in consumer and industrial products, posing significant public health risks and contributing to the onset of endocrine and reproductive related diseases. Parabens are a suspected class of EDCs (Nowak et al. 2018). These chemicals are alkyl esters of para-hydroxybenzoic acid, which are extensively used as preservatives or antimicrobial agents in various cosmetics, pharmaceuticals, and food products, and include compounds such as methyl-, butyl-, propyl-, and benzyl paraben (Mitra et al. 2021). They are present in more than 22,000 cosmetic formulations, either as mixtures or individual constituents, with concentrations up to 0.8%. Among parabens, propyl paraben has been shown to impair reproductive function, raising concerns about its safety (Oishi 2002b; Gal et al. 2019; Jiao et al. 2021). Benzyl paraben (BzP) and butyl paraben (BuP) remain understudied, and their ability to perturb hormone-dependent processes is not well understood (**Figure 1**).

**Figure 1.**
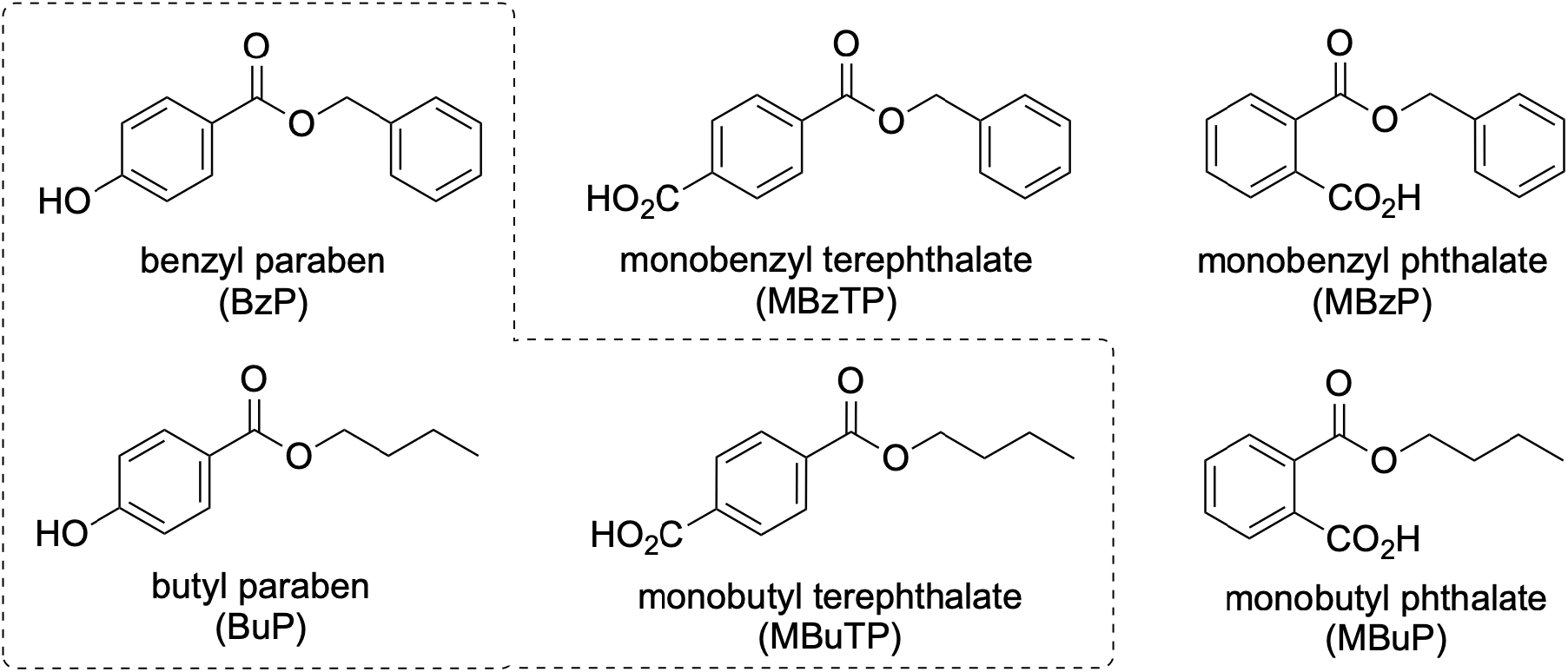
Chemical structures of benzyl paraben (BzP), butyl paraben (BuP), monobenzyl terephthalate (MBzTP), monobutyl terephthalate (MBuTP), monobenzyl phthalate (MBzP), and monobutyl phthalate (MBP). Dotted box indicates chemicals experimentally investigated in this study. The other three compounds are structurally similar for comparison.

Widespread human exposure to parabens from consumer products challenges the adequacy of current regulatory frameworks. Current regulatory guidelines establish acceptable daily intake (ADI) of 10 mg/kg body weight per day (bw/d) for methyl and ethyl parabens and 0.1 mg/kg for propyl paraben; no ADI exist for BzP or BuP (Opinion of the Scientific Committee on Consumer Products on the safety evaluation of parabens 2005; Boberg et al. 2010; Ihn et al. 2026). The global average for grouped estimated daily intakes of the most common parabens (methyl, ethyl, propyl, and butylparaben) were 8.00, 1.06, 2.50, and 0.27 μg/kg bw/day, respectively, based on urinary biomarker levels (Liao et al. 2025), indicating routine, multi-route exposure. Some populations have significantly higher exposures. In children, for example, daily intake values of paraben constituents through liquid medication can reach maximum estimates of 740 μg/kg for methylparaben, 640 μg/kg for propylparaben, and 440 for μg/kg for butylparaben (Ihn et al. 2026). This concern is further augmented by growing biomonitoring evidence in urine and tissues, including breast tissue, menstrual blood, placental tissue, and ovarian follicular fluid (Barr et al. 2012; Vela-Soria et al. 2015; Jiménez-Díaz et al. 2016; Bellavia et al. 2023; Tian et al. 2023; Shulhai et al. 2024). Studies also indicate the widespread occurrence of parabens in various environmental matrices (Núñez et al. 2008; Wang et al. 2012; Chen et al. 2021; Zhang et al. 2021). The combination of widespread exposure and the presence of parabens in key reproductive tissues highlights the potential for these chemicals to impair reproductive function (Pulcastro and Ziv-Gal 2024).

Consistent with these findings, epidemiological and rodent studies have linked internal paraben exposure to endocrine disruption, including reduced dehydroepiandrosterone sulfate (DHEA-S), 17-hydroxyprogesterone, and follicle stimulating hormone (FSH) concentrations in female infants, lower estradiol concentrations and estradiol-to-progesterone ratios during pregnancy, and reduced umbilical cord blood DHEA levels compared to populations with lower paraben exposure (Kolatorova et al. 2018; Aker et al. 2019; Jensen et al. 2021). Prenatal exposure to methyl- and propylparaben has also been associated with lower birth weight and impaired neonatal growth parameters in female, but not male, offspring, suggesting sex-specific developmental effects (Chang et al. 2019; Pacyga et al. 2022). In mice, butylparaben and propylparaben altered ovarian steroidogenesis and follicle numbers in neonatal mice (Ahn et al. 2012). In adults, chronic low dose propylparaben exposure altered estrus cyclicity and hormone levels and *in vitro* exposure to propylparaben inhibited ovarian antral follicle growth and increased production of testosterone and estradiol (Gal et al. 2019; Yan et al. 2022). Collectively, these findings suggest that internal paraben exposure may adversely affect female reproductive health, even at exposure levels permitted under current regulatory guidelines.

Molecular evidence also suggests that parabens are endocrine disruptors, although previous studies have focused mostly on propylparaben and estrogenic activity (Isaac et al. 2025). Further rationale for investigating BzP and BuP comes from their structural similarity to phthalate and terephthalate metabolites, including monobenzyl phthalate (MBzP), monobenzyl terephthalate (MBzTP), monobutyl phthalate (MBP), and monobutyl terephthalate (MBuTP) (**Figure 1**). Terephthalates are of particular interest because they are rising replacements for phthalates and feature the same para-substituted benzene ring structure as parabens. Monoester phthalate metabolites and the most common terephthalate metabolite, mono-2-ethylhexyl terephthalate have been shown to disrupt ovarian endocrine function (Hannon et al. 2015; Rowdhwal and Chen 2018; Warner et al. 2021; Potts et al. 2025), but little is known about other terephthalate metabolites despite their detection in human biomonitoring and plastic extracts (Nayebare et al. 2018; Monclús et al. 2025). The structural similarities raise the question of whether BzP and BuP share toxicological mechanisms with their structural analogues, such as activation of peroxisome proliferator activated receptors (PPARs) (Meling et al. 2022; Alahmadi et al. 2024).

In this study, we utilized an *in vitro* model culturing antral follicles from CD-1 mice to assess the ovarian toxicity of BzP and BuP. Based on interesting findings with BuP, we selected MBuTP for follow up experiments and used all 6 compounds (Figure 1) for in silico comparison. The ovary is highly conserved in mammals, thus providing translatability to human health (Hirshfield 1991). This study focuses on the primary ovarian functions of steroid hormone production (steroidogenesis) and follicle development (folliculogenesis) to provide insight into the potential of BzP, BuP, and MBuTP to act as an ovarian toxicants and endocrine disruptors at human-relevant doses.

## 2. Methods

### 2.1 Chemicals

BzP was obtained from LGC Standards (TRC-B288575, ≥ 95% purity), BuP from Fisher Scientific (AC403571000, ≥ 99% purity), and MBuTP from Chemdiv (1136-0002, ≥ 98% purity). For the *in vitro* studies, follicles were exposed to vehicle control (dimethyl sulfoxide, DMSO, Sigma Aldrich D2650, ≥ 99.7% purity) or 0.1, 1, 10, or 100 µg/mL of BzP, BuP or MBuTP. The doses were chosen for environmental relevance and for comparison to previous studies with parabens in the ovary (Gal et al. 2019). The 0.1-10 µg/mL are representative of everyday human exposure from parabens based on previous biomonitoring studies (Ihn et al. 2026); the 100 µg/mL concentrations are high doses for comparison to previous studies with other EDCs and for providing mechanistic insight. Treatment solutions of 0.13, 1.33, 13.3, and 133.3 mg/mL of BzP, BuP or MBuTP were prepared by diluting each chemical in DMSO such that the same volume of the treatment solution (0.75 µL) could be added per mL of culture media to maintain consistent DMSO concentration of 0.075%.

### 2.2 Animals

Young adult female CD-1 mice (23 days old) were purchased from Charles River (Wilmington, MA). They were acclimatized in a controlled environment at the Rutgers University Newark Animal Facility for at least 7 days, with a 12-hour light-dark cycle, 22 ± 1 °C temperature, and ad libitum access to food and water. All experiments were performed with Rutgers Institutional Animal Care and Use Committee (IACUC) approval (PROTO202100043).

### 2.3 Antral follicle culture

Untreated adult female CD-1 mice were euthanized at 32–38 days of age. Antral follicles were manually isolated from the ovaries based on relative follicle size (200–400 µm). Interstitial tissue was cleaned using watchmaker forceps. Follicles were obtained from 3-5 mice and were individually plated in 96 well culture plates with 10-12 follicles per group. The different treatment groups (DMSO or 0.1–100 µg/mL) were supplemented in minimum essential medium alpha (Gibco) containing 1% insulin-transferrin-selenium (10 μg/ml insulin, 5.5 μg/ml transferrin, 5 ng/ml sodium selenite, Sigma-Aldrich), 100 U/ml penicillin (Sigma-Aldrich), 100 μg/ml streptomycin (Sigma-Aldrich), 5 IU/mL human recombinant follicle-stimulating hormone (Scripps Laboratories), and 5% fetal bovine serum (Thermo Fisher). Stock concentrations of BzP, BuP and MBuTP were prepared for each dose (0.1, 1, 10, 100 µg/mL), and 0.75 µL of stock was used per 1 mL of supplemented medium. Follicles were cultured for 24 hours for cytotoxicity analysis for both BzP and BuP or 96 hrs for all other analyses in an incubator supplying 5% CO2 at 37 °C.

### 2.4 Lactate Dehydrogenase (LDH) Colorimetric Activity assay analysis

To assess cytotoxic activity, a lactate dehydrogenase (LDH) colorimetric activity assay was used, specifically the CyQUANT™ LDH Cytotoxicity Assay Kit from Thermo Fisher. To analyze cytotoxic activity, media was analyzed after 24 hours of culture. The assay was performed according to the manufacturer’s instructions and read with an Agilent Biotek Epoch microplate reader. Absorbance was measured at 490 nm and 680 nm. Percent cytotoxicity was calculated as the ratio of the treatment induced LDH activity to the baseline LDH activity.

### 2.5 Follicle growth analysis

Follicle growth was assessed every 24 hours for the 96 hour culture period by measuring follicle diameters on perpendicular axes with an inverted microscope equipped with a calibrated ocular micrometer. The diameters of each follicle were averaged within the treatment group for each 24 hr interval, and the average values were divided by the initial average measurement at 0 hr of each of the respective treatment groups to calculate the percent change in follicle diameter over time. The percent change in antral follicle diameter over time was used for statistical analysis.

### 2.6 Hormone assays

After the 96 hour follicle cultures were completed, media were pooled by treatment group and frozen at -80 °C until further analysis. Hormone analysis was performed on media from *in vitro* experiments via enzyme linked immunosorbent assay (ELISA). Kits for estradiol, testosterone, progesterone, androstenedione, and pregnenolone were purchased from DRG International, Inc. (Mountainside, New Jersey) and used according to the manufacturer’s instructions. Samples were run in duplicate. Samples were diluted as needed to match the dynamic range of each ELISA kit and read with an Agilent Biotek Epoch microplate reader. The detection ranges were 0.0083–16 ng/ml for testosterone, 0–40 ng/ml for progesterone, 10.6–2000 pg/ml for estradiol, and 0–10 ng/ml for androstenedione. Hormone concentrations were interpolated using GraphPad Prism software. The mean values of each sample were used for statistical analysis. Interassay and intrassay coefficients of variability were ≤ 15% for all ELISAs.

### 2.7 Gene expression analysis

After 96 hours of culture, follicles from each treatment group were combined and stored at −80 °C for later analysis. Total RNA was extracted from the follicles using the Qiagen RNeasy Micro Kit according to the manufacturer’s protocol. RNA concentrations were quantified using a Thermo Scientific Nanodrop One C Spectrophotometer. cDNA was synthesized from 100 ng RNA using an iScript RT kit (Bio-Rad Laboratories, Inc., Hercules, California). The quantitative polymerase chain reaction (qPCR) reactions were run in duplicate with 1.67 ng cDNA and 7.5 pmol gene-specific primers (Integrated DNA Technologies, Inc. Coralville, Iowa) for a final reaction volume of 10 µl. Expression levels of aromatase (*Cyp19a1*), 17a-hydroxylase (*Cyp17a1*), 17b-hydroxysteroid dehydrogenase 1 (*Hsd17b1*), cytochrome P450 11A1 (*Cyp11a1*), 3-beta- hydroxysteroid dehydrogenase 1 (*Hsd3b1*), steroidogenic acute regulatory protein (*Star*), Beta actin (*ActB),* caspase 3 (*Casp3)*, caspase 8 (*Casp8*), Bcl2-like 10 *(Bcl2l10*), B cell leukemia/lymphoma 2 *(Bcl2*), BH3 interacting domain death agonist (*Bid*), Bcl2-associated X protein (*Bax*), Bcl2-associated agonist of cell death (*Bad*), antigen identified by monoclonal antibody Ki 67 (*Ki67),* cyclin dependent kinase inhibitor 2A (*Cdkn2a*), cyclin dependent kinase inhibitor 1A (*Cdkn1a*), cyclin dependent kinase inhibitor 2B (*Cdkn2b)*, cyclin dependent kinase inhibitor 2D (*Cdkn2d*), cyclin A2 (*Ccna2*), cyclin dependent kinase 4 (*Cdk4*), cyclin dependent kinase 2 (*Cdk2*), cyclin D2 (*Ccnd2*), cyclin D1 (*Ccnd1*), perilipin-1 (*Plin1*), angiopoietin-like 4 (*Angptl4*), fatty acid binding protein 4 (*Fabp4*), CD 36 molecule (*Cd36*) (**Table S1**). *ActB* was used as a reference gene for normalization for *in vitro* samples (Hannon et al. 2015). Following the completion of qPCR, the Pfaffl method was used to obtain relative fold changes in comparison to controls (Pfaffl 2001).

### 2.8 Molecular docking

The structure of the PPARγ (PDB ID: pdb_00001FM9) receptor was downloaded from the Research Collaboratory for Structural Bioinformatics Protein Data Bank (RCSB PDB) database (Gampe et al. 2000). PyMOL was used to remove any ion, water molecules, ligands and receptors that were not of interest to the experiment (Schrödinger, LLC 2015). The molecular docking was performed on CB-DOCK2, which is web based docking server that uses AutoDock as the background to generate the free-energy or bind affinity and the binding poses between the ligand and the receptors (Liu et al. 2022; Yang et al. 2022). Once the file was clean it was saved and uploaded to CB-DOCK2. The ligands were uploaded to CB-DOCK2 as SMILES codes. The program generated a vina score and determined the binding affinity between the ligand and receptor. A ligand/receptor with a vina score that is lower than -7.0 Kcal/mol is considered to have a strong/moderate binding affinity (Alsedfy et al. 2024). Lower (more negative) vina scores indicate stronger binding affinity. PDB files for pairing were uploaded to Pymol to visualize the ligand within the binding pocket. Pioglitazone was used as the positive control for PPARγ.

### 2.9 Statistical analysis

Data were expressed as the mean ± standard error of the mean (SEM). Missing values from undetectable samples were imputed as the minimum detectable value divided by the square root of 2 for ELISAs and as the maximum detectable cycle value (36) for qPCR. Data were analyzed by comparing treatment groups to control using IBM SPSS version 29 software (SPSS Inc., Chicago, IL, USA). All data were continuous and assessed for normal distribution by Shapiro-Wilk analysis. If data met assumptions of normal distribution and homogeneity of variance, data were analyzed by one-way analysis of variance (ANOVA) followed by Tukey HSD or Dunnett 2- sided *post-hoc* comparisons. However, if data met assumptions of normal distributions, but not homogeneity of variance, data were analyzed by ANOVA followed by Games-Howell or Dunnett’s T3 *post-hoc* comparisons. If data were not normally distributed or presented as percentages, the independent sample Mann Whitney U non-parametric tests were performed. For all comparisons, statistical significance was determined by p-value ≤ 0.05. If p-values were greater than 0.05, but less than 0.10, data were considered to exhibit a trend towards significance.

## 3. Results

### 3.1 Effects of BzP and BuP on follicle growth in vitro

To assess the effects of BzP and BuP on follicle growth, isolated antral follicles were cultured in media containing vehicle control (DMSO) or various doses of BzP or BuP for 96 hours. BzP and BuP treatment at 100 µg/mL significantly decreased follicle growth at every time point compared to control (**Figure 2**). BuP at 10 and 1 µg/mL significantly decreased follicle growth at 72, and 96 hours respectively compared to control. BzP at 0.1 µg/mL also trended towards a significant decrease at 96 hours compared to control.

**Figure 2:**
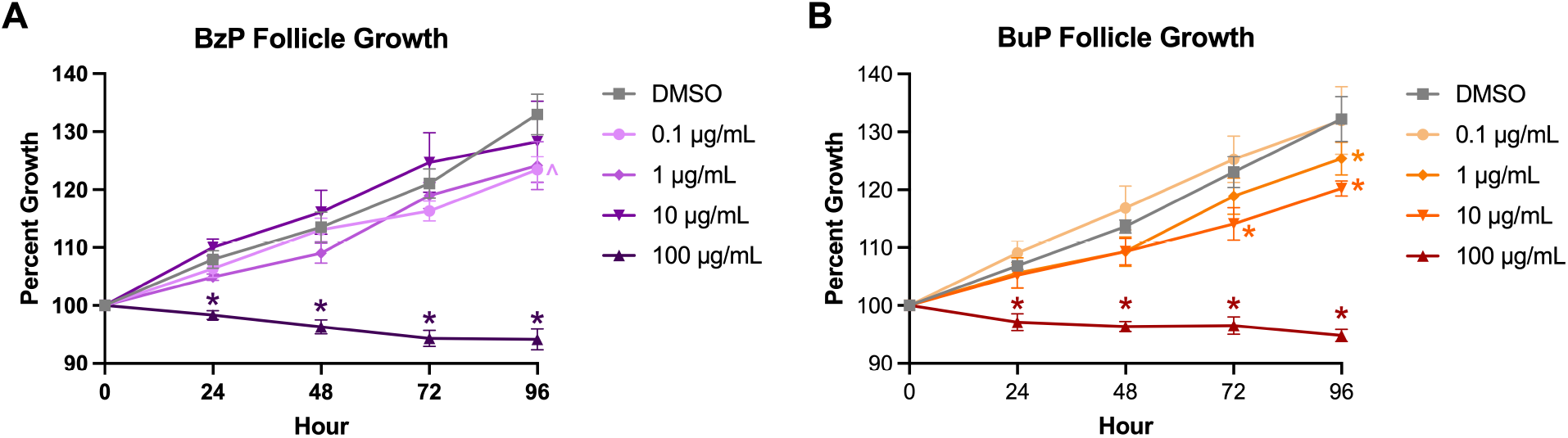
Effects of 96 hr *in vitro* BzP (A) and BuP (B) treatment on antral follicle growth. Follicle growth was measured every 24 hours for 96 hours. Graphs represent means ± SEM from 6-7 independent experiments per treatment group. Asterisks (∗) indicate significant differences from the control (p ≤ 0.05) and carets (^) indicates a trend toward significance (p ≤ 0.10).

After observing that the highest dose of parabens resulted in consistent follicle death, we suspected that cytotoxicity was occurring. We assessed cytotoxicity at 24 hours because the 100 µg/mL treatment group showed decreased average follicle size at this first time point. LDH activity was used to measure cytotoxicity, with values expressed as percent change relative to the control; values near 0% indicate little to no cytotoxicity. At 24 hours, the 100 µg/mL dose of both BzP and BuP showed the greatest LDH-associated cytotoxicity (**Figure 3**).

**Figure 3:**
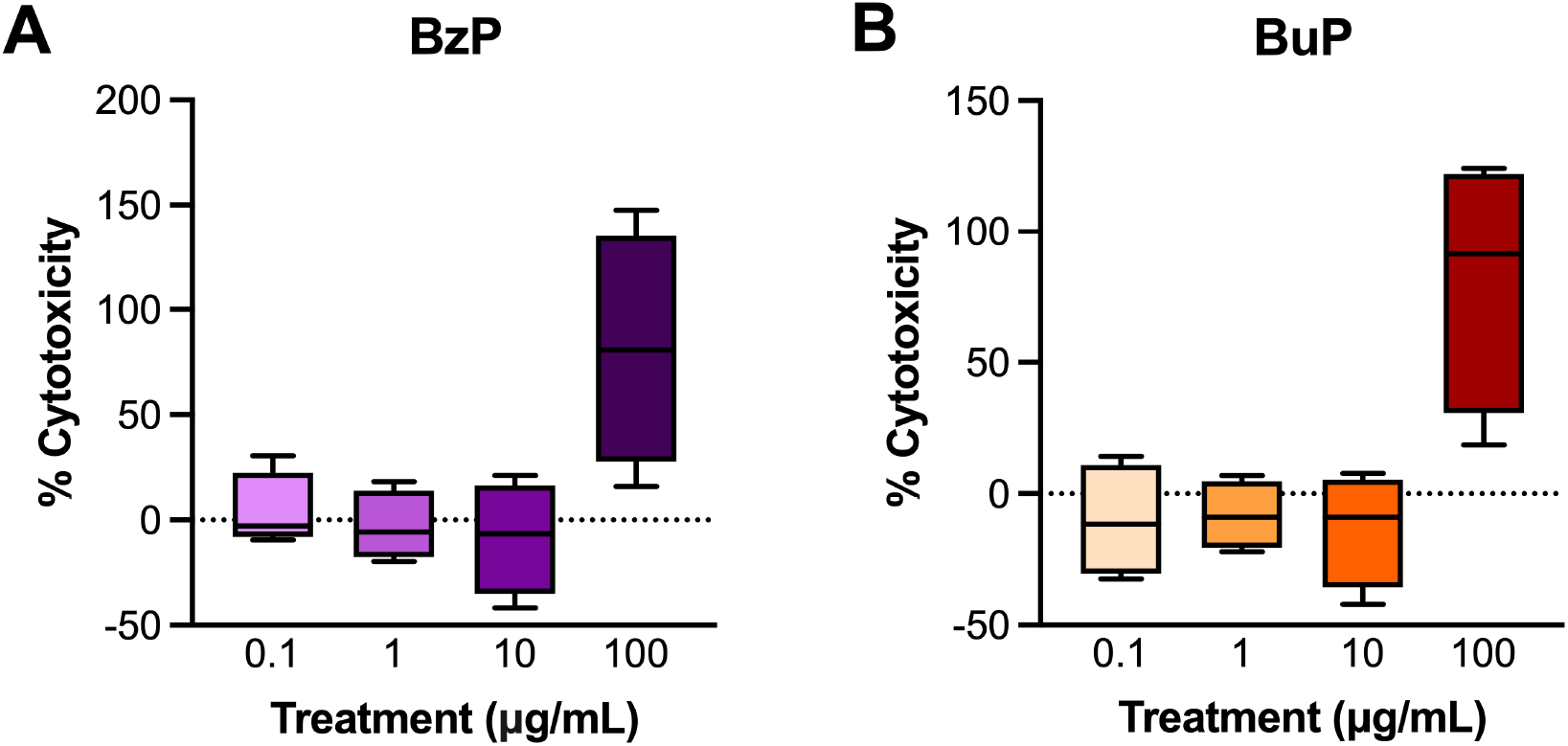
Effects of BzP (A) and BuP (B) on cytotoxic activity after 24 hours of antral follicle culture. Cytotoxicity was quantified via LDH release and expressed as a percentage relative to control; values near 0% indicate little to no cytotoxic activity.

### 3.2 Effects of BzP and BuP exposure on gene expression of cell cycle and apoptosis regulators in vitro

Although the 100 μg/mL BzP and BuP treatment groups were not suitable for qPCR analysis, gene expression analysis remained feasible at the lower doses (0.1, 1, and 10 μg/mL). The selected cell cycle regulators and apoptotic markers were chosen based on pathways previously shown to be affected by phthalate exposure. No significant effects on cell cycle regulators were observed at 0.1, 1, and 10 μg/mL of BzP treatment compared to control (**Figure 4A**). *Cdkn2a* and *Cdkn2d* were significantly downregulated by 0.1 μg/mL BuP compared to control and borderline downregulated at 1 μg/mL BuP (**Figure 4B).** *Cdkn2b* was borderline upregulated at 10 μg/mL BuP expression levels compared to control. *Cdk2* was significantly downregulated by 0.1 and 10 μg/mL BuP compared to control.

**Figure 4:**
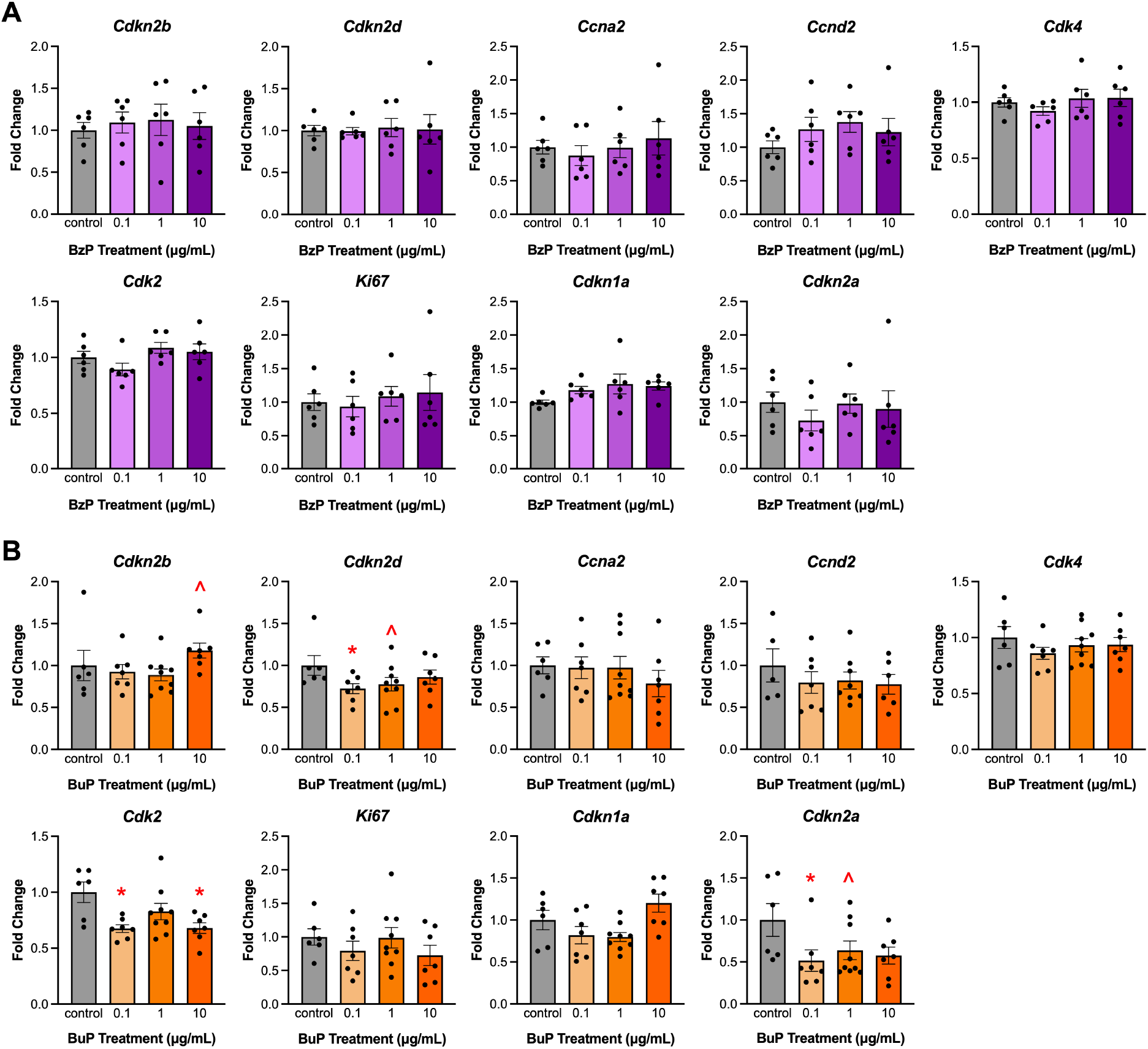
Effects of *in vitro* BzP (A) and BuP (B) on cell cycle regulator gene expression in antral follicles expressed as fold change compared to control. All gene expression is relative to the housekeeper gene *B-actin*. Graphs represent means ± SEM from 6-7 experiments per treatment group. Asterisks (∗) indicate significant differences from the control (p ≤ 0.05) and carets (^) indicates a trend toward significance (p ≤ 0.10).

BzP treatment did not alter expression of apoptosis regulators compared to control (**Figure S1A**). Expression of the pro-apoptotic regulator *Bad* was significantly reduced at 0.1 μg/mL BuP relative to control (**Figure S1B**).

### 3.3 Effects of BzP and BuP on sex steroid hormone levels and steroidogenic enzyme gene expression in vitro

96 hrs of BzP exposure at 0.1 µg/ml borderline increased testosterone and 10 µg/mL significantly increased testosterone compared to control (**Figure 5A**). BzP exposure at 1 µg/mL significantly increased estradiol and 10 µg/mL borderline increased estradiol compared to control. BuP did not alter hormone levels at 0.1-10 µg/ml (**Figure 5B**), but expression of *Cyp17a1* was significantly reduced compared to control in all treatment groups (**Figure 5D**).

**Figure 5:**
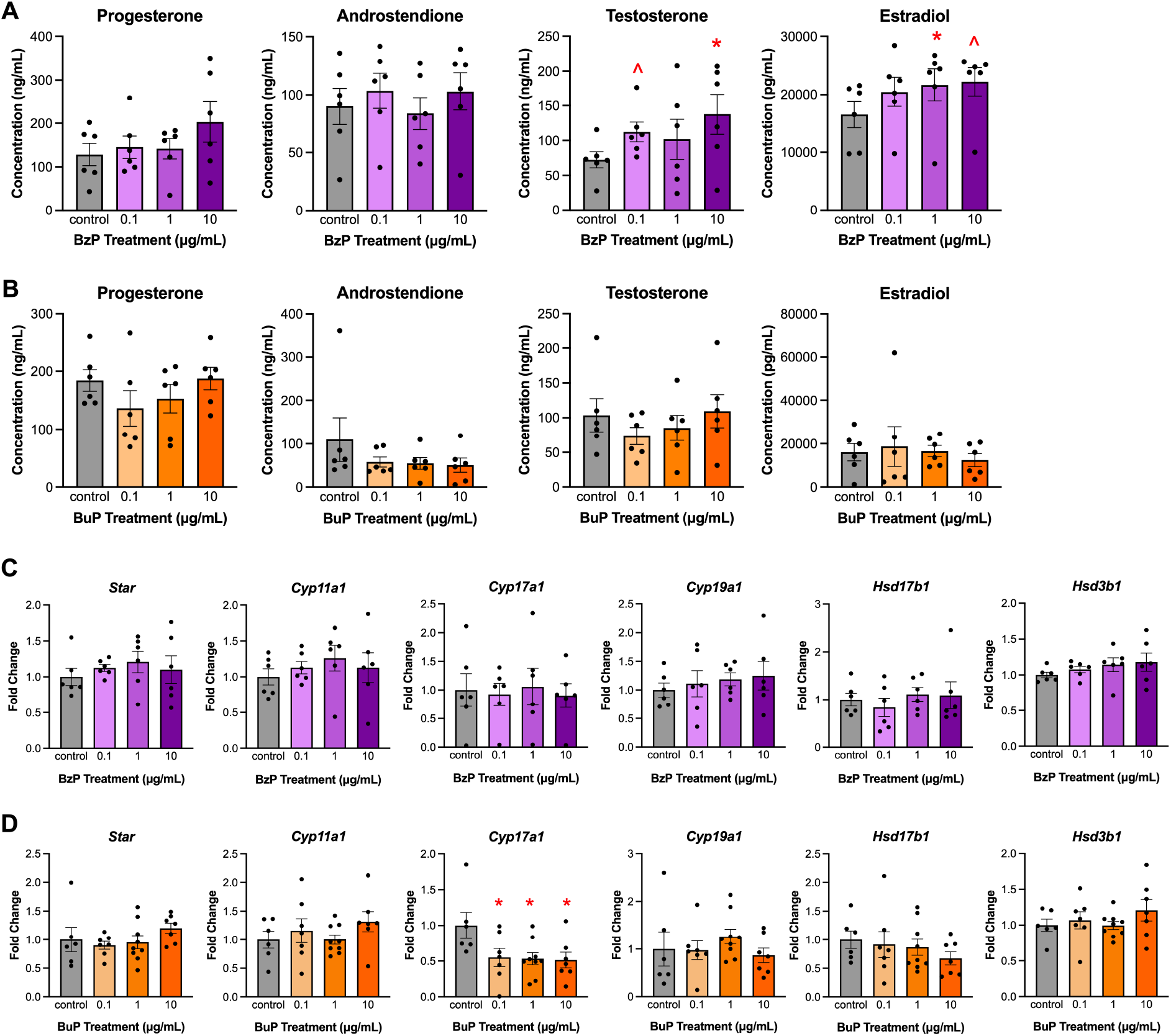
Effects of 96 hr *in vitro* BzP and BuP treatment on sex steroid hormone levels (A, B) and steroidogenic enzyme gene expression (C, D) in antral follicles. All gene expression is relative to the housekeeper gene *B-actin*. Graphs represent means ± SEM from 6-7 experiments per treatment group. Asterisks (∗) indicate significant differences from the control (p ≤ 0.05) and carets (^) indicates a trend toward significance (p ≤ 0.10).

### 3.4 Effects of BzP and BuP on genes downstream of PPARs

To assess PPAR activation by parabens, we measured expression of genes under direct PPAR control known to be activated by phthalates. BzP and BuP exposure at 10 μg/mL significantly upregulated only *Fabp4* expression compared to the control (**Figure 6**).

**Figure 6:**
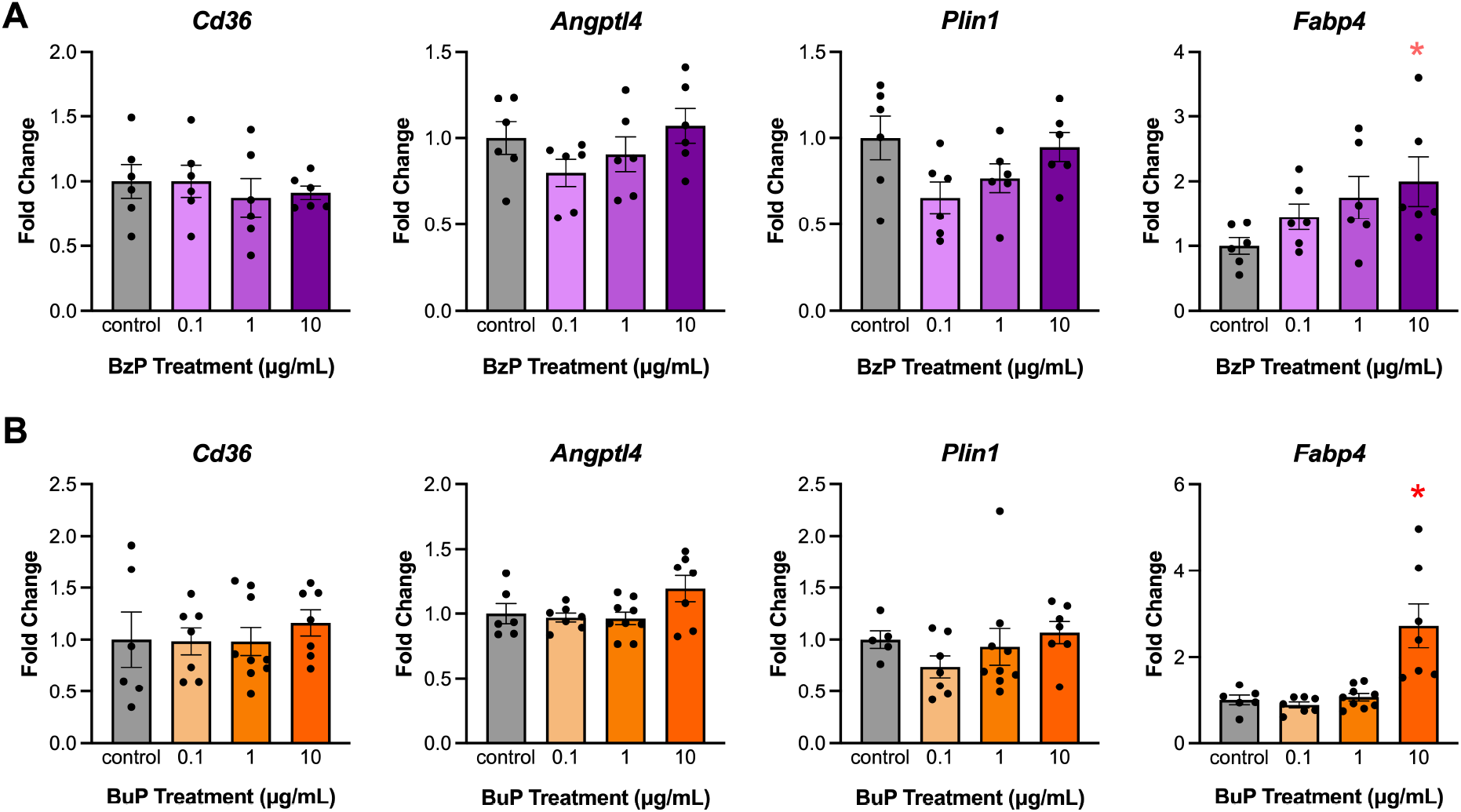
Effects of 96 hr *in vitro* BzP (A) and BuP (B) treatment on peroxisome proliferator-activated receptor (PPAR) controlled genes in antral follicles. All gene expression is relative to the housekeeper gene *B-actin*. Graphs represent means ± SEM from 6-7 experiments per treatment group. Asterisks (∗) indicate significant differences from the control (p ≤ 0.05).

### 3.5 Effects of MBuTP on follicle growth, cell cycle regulator gene expression, steroidogenic enzyme gene expression, and sex steroid hormone levels

MBuTP treatment at 100 μg/mL significantly reduced follicle growth at 24, 72, and 96 hours (**Figure 7**). MBuTP treatment at 0.1 and 10 μg/mL significantly reduced follicle growth at 72 and 96 hours. MBuTP at 1 μg/mL significantly reduced follicle growth at only 96 hours compared to control.

**Figure 7:**
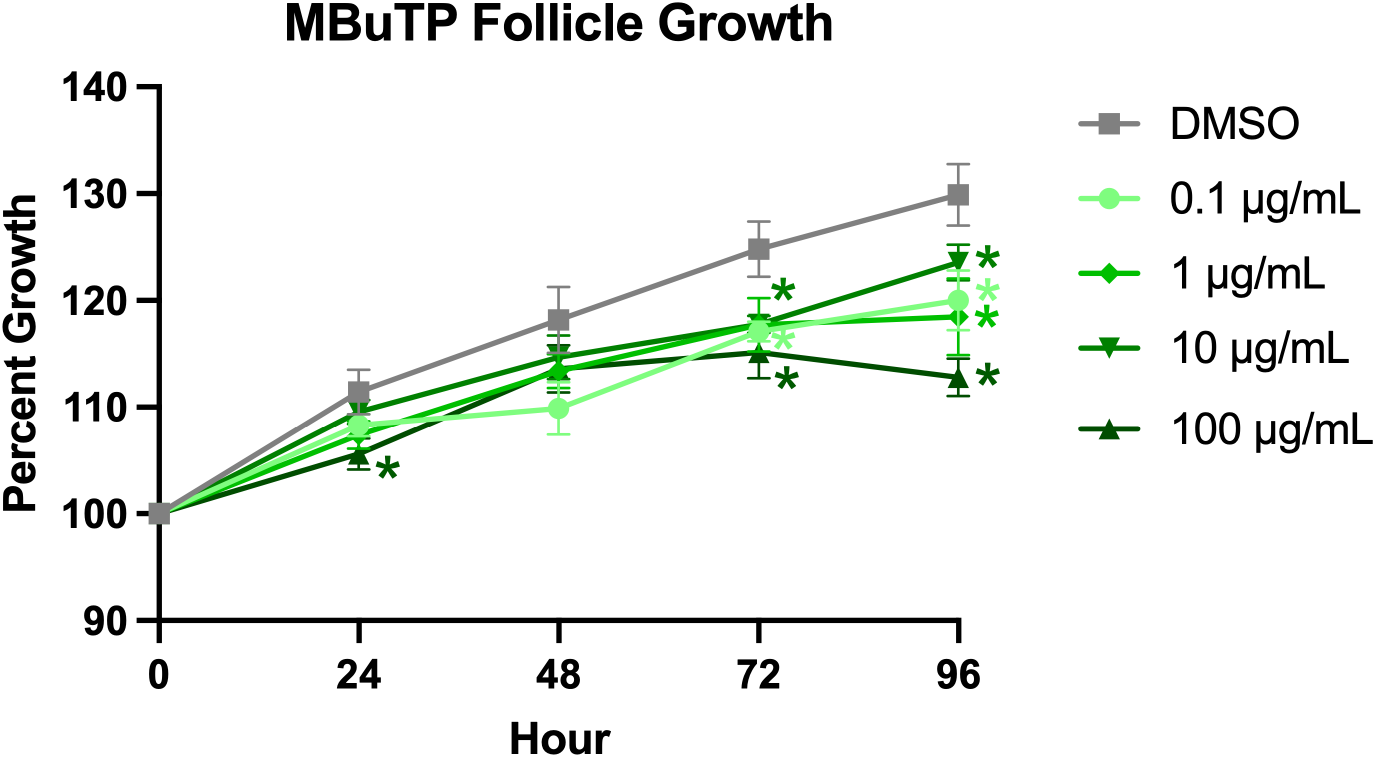
Effects of *in vitro* MBuTP treatment on antral follicle growth. Follicle growth was measured every 24 hours for 96 hours. Graphs represent means ± SEM from 5 independent experiments per treatment group. Asterisks (∗) indicate significant differences from the control (p ≤ 0.05).

MBuTP exposure at 100 μg/mL significantly upregulated expression of cell cycle inhibitor *Cdkn1a* compared to control (**Figure S2**). Exposure to MBuTP did not significantly alter steroidogenic enzyme gene expression (**Figure S3A**). 100 and 10 μg/mL MBuTP borderline increased estradiol levels (**Figure S3B)**.

### 3.6 Effects of MBuTP on gene expression of peroxisome proliferator activated receptors in vitro

To evaluate effects of MBuTP exposure on PPAR-mediated pathways, gene expression analyses were conducted after 96 hours. Genes downstream of PPARα and/or PPARγ were selected based on established links with phthalate activity (Meling et al. 2022). MBuTP at 100 μg/mL significantly upregulated expression levels of *Cd36*, *Angptl4*, *Plin1*, and *Fabp4* compared to the control (**Figure 8**).

**Figure 8:**
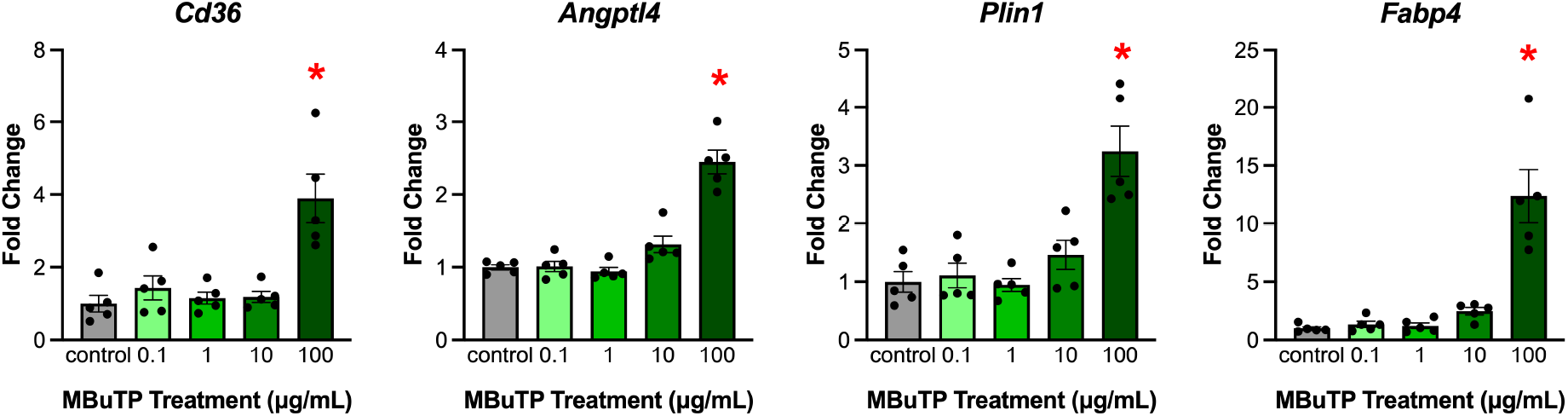
Effects of 96 hr *in vitro* MBuTP treatment on peroxisome proliferator-activated receptor (PPAR) controlled genes in antral follicles. All gene expression is relative to the housekeeper gene *B-actin*. Graphs represent means ± SEM from 5 experiments per treatment group. Asterisks (∗) indicate significant differences from the control (p ≤ 0.05).

### 3.7 Docking of BzP, BuP, and their monoester phthalate and terephthalate analogs to PPARγ

*Fabp4* is a directly downstream target of PPAR*γ*, so we selected PPAR*γ* for molecular docking simulations using all six compounds of interest (**Figure 1**). Vina scores represent predicted binding energy in Kcal/mol, with more negative values as stronger binders. In general, benzyl substituent compounds had stronger predicted binding than butyl substituent compounds (**Figure 9A**). The scores for each set of three compounds with the same side chain were similar. The benzyl compounds were predicted to bind with similar affinity to the positive control ligand, the pharmaceutical PPAR*γ* activator pioglitazone. When the binding interactions were visualized, the three compounds of each side chain closely overlay, although there are some side chain orientation variations.

**Figure 9:**
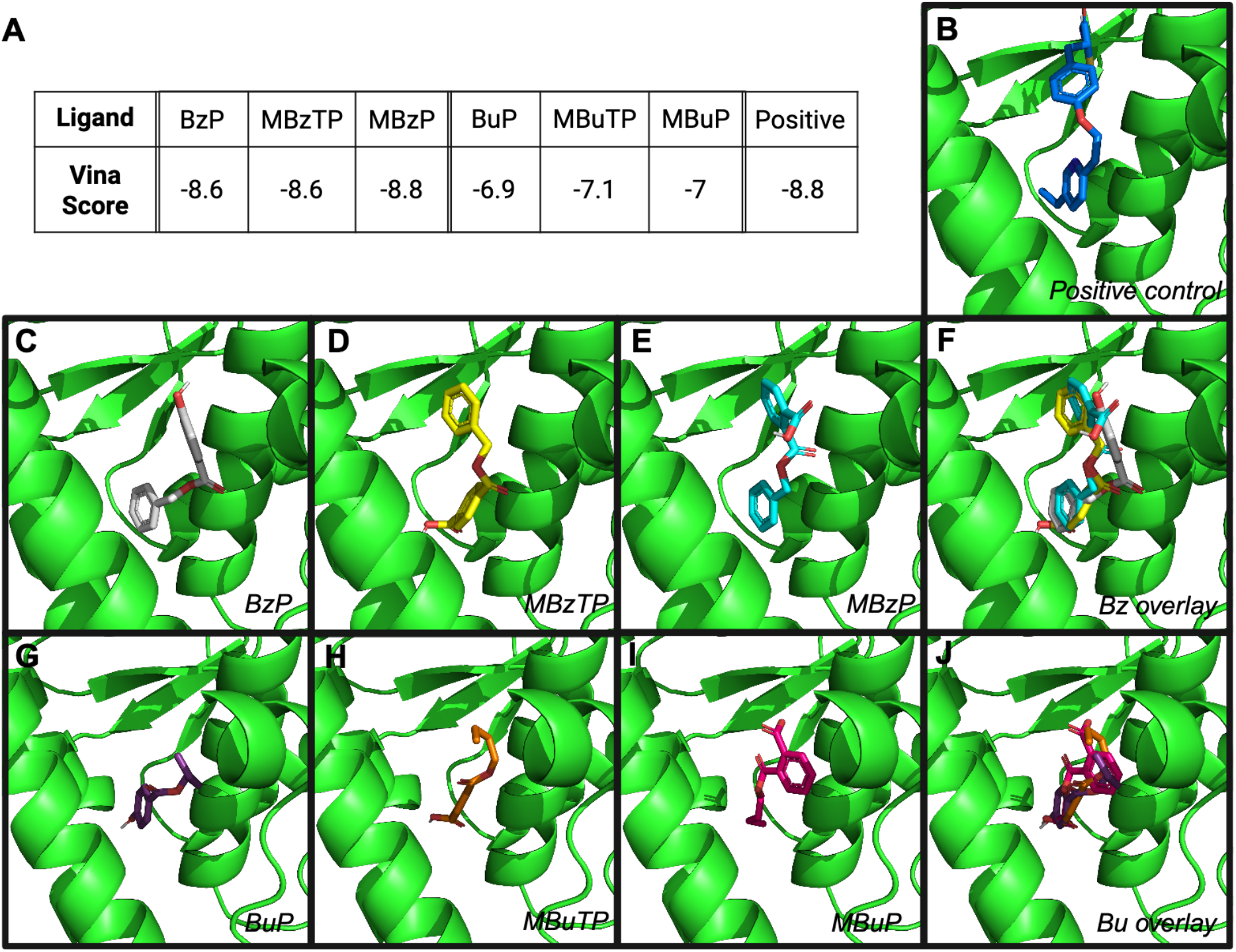
Computational modeling of the interaction between PPARγ and benzyl or butyl paraben, monoester terephthalate, or monoester phthalate with pioglitazone as positive control. Vina scores in Kcal/mol between PPARγ and each compound (A). Within the PPARγ binding pocket, pioglitazone (B), BzP (C), MBzTP (D), MBzP (E), BuP (F), MBuTP (G), MBuP (H), overlay of BzP, MBzTP, and MBzP (I), overlay of BuP, MBuTP, and BuP (J)

## 4. Discussion

Parabens are esters of para-hydroxybenzoic acid, with a substituted ester opposite a hydroxy group on the aromatic ring. Terephthalates, rising replacements for phthalates as plasticizers, are para-substituted isomers of phthalates, meaning they have the same ring substitution pattern as parabens. Terephthalate biological metabolites are monoesters, with a substituted ester opposite a carboxylic acid, such that parabens and terephthalate metabolites have only one functional group different in their structures (**Figure 1**). With our history of studying phthalates and recent interest in phthalate replacements, this similarity caught our attention and frames our interest in parabens. Terephthalates such as di-2-ethylhexyl terephthalate (DEHTP) and metabolites are widely detected in biomonitoring at higher concentrations than phthalates (Potts et al. 2026). Although there is little biomonitoring of shorter chain terephthalates, we know they are used in consumer products from plastic sampling studies (Monclús et al. 2025). Para-substituted plasticizers are promoted as safer alternatives to traditional phthalates, but their similarities to other chemicals have so far been overlooked. To our knowledge, this is the first study to analyze the similarities between terephthalates and parabens in toxicity and molecular mechanisms of action. Parabens are widely used in consumer products and detected in human tissues. Previous studies suggest that BzP and BuP have disruptive effects on the reproductive system and may be endocrine disruptors, but little is known about their ovarian toxicity at environmentally relevant levels (Oishi 2002a; Darbre et al. 2003; Maske et al. 2018). In this study, we exposed antral follicles from young adult mice to BzP and BuP in culture. Antral follicles are the most mature ovarian follicle type. Each contains an oocyte maturing toward ovulation (or atresia) and granulosa and theca cells that produce hormones. We measured follicle size every day of culture, as growth is an important metric for folliculogenesis, the process of follicle development. Most phthalates are not or minimally cytotoxic up to 100 µg/mL, which is a supraphysiological concentration (Meling et al. 2022). Thus, reduced growth at exposures of up to 100 µg/mL typically suggests cell cycle impairment or increased cellular atresia (Zhou and Flaws 2017). However, follicles cultured with 100 µg/mL of BzP or BuP had no growth and appeared dead by the end of the 96 hr culture period, suggesting higher cytotoxicity compared to phthalates. Propylparaben similarly inhibited follicle growth in a previous study (Gal et al. 2019).

We confirmed cytotoxicity after 24 hours of culture, with no cytotoxicity observed at 10 µg/mL or below. Although 100 µg/mL is likely supraphysiological for parabens as well, paraben levels in young children from medications is orders of magnitude higher than average adult exposure, with the Korean National Environmental Health Survey reporting 95%ile urine concentrations of > 5 µg/mL creatinine corrected and US National Health and Nutrition Examination Survey reporting > 2 µg/mL creatinine corrected for methyl paraben in children 3-5 years old (Hong et al. 2021). Our results are in agreement with previous cytotoxicity studies in human epidermal and dermal cell lines that suggest increasing paraben cytotoxicity with increasing chain length (Chrz et al. 2020; Janiga-MacNelly et al. 2025).

BuP slowed follicle growth at 1 and 10 µg/mL, which is consistent with disruption of cell cycle regulators in the gene expression results. Phthalates typically inhibit cell cycle promoters and increase expression of inhibitors, resulting in cell cycle arrest, especially at the G1/S transition (Craig et al. 2013; Zhou and Flaws 2017). BuP exposure decreased expression of both inhibitors and promoters, except for a trending increase in one inhibitor, *Cdkn2b*. *Cdkn2b* has been consistent upregulated in previous phthalate and terephthalate studies (Meling et al. 2020; Warner et al. 2021; Potts et al. 2025). Similarly, *Cdk2* was downregulated by BuP exposure, consistent with phthalate and terephthalate exposures (Meling et al. 2020; Potts et al. 2025) impacting G1/S phase transition. Interestingly, MBuTP exposure decreased follicle growth but only upregulated the inhibitor *Cdkn1a*, which is consistent with previous studies of the same side chain phthalate, dibutyl phthalate (Craig et al. 2013; Rasmussen et al. 2017).

Peroxisome proliferator-activated receptors are ligand-activated nuclear transcription factors that regulate lipid metabolism, glucose homeostasis, and cellular differentiation; within the ovary, they modulate follicular development, steroidogenesis, and granulosa cell function by controlling genes involved in lipid utilization and hormone synthesis (Chinetti et al. 2000). Of note, *Fabp4* was significantly upregulated by both BzP and BuP at 10 µg/mL. *Fabp4* is highly enriched in adipocytes, and commonly used as a marker of PPARγ activation, a nuclear receptor that, upon activation by fatty acid ligands, promotes the transcription of genes involved in lipid uptake and storage (Komar 2005; Fang et al. 2016). A previous study on phthalate monoesters in CD-1 mouse granulosa cells reported that monobenzyl phthalate and monobutyl phthalate significantly upregulated *Fabp4* expression (Meling et al. 2022).

We measured expression of three other genes under PPAR transcriptional control and disrupted by phthalates in previous ovary work, *Cd36*, *Angptl4*, and *Plin*1 (Alahmadi et al. 2024). All three were activated by MBuTP like classic phthalates, whereas none were activated by paraben exposure, suggesting that parabens may be weaker or more selective activators. Our modeling of ligand binding pocket-PPARγ interactions suggest that all six benzyl and butyl ligands fit similarly, but the phthalates and terephthalates had slightly stronger predicted interactions compared to the parabens. Alternatively, it is possible that paraben *Fabp4* activation may be due to a non-PPAR mechanism.

This study provides insight into the mechanisms of paraben toxicity at doses that are relevant to human exposure in a human-translational *ex vivo* model that is sensitive to endocrine disruption. We observed changes in hormone levels from exposure to benzyl paraben, whereas butyl paraben had similar effects as phthalates, especially dibutyl phthalate, on the cell cycle. Most of the cell cycle regulator impacts suggested non-monotonic dose responses characteristic of phthalates and endocrine disruption. However, there are several limitations to this study. Gene expression does not necessarily translate to protein levels, and we only measured a small selection of cell cycle regulators and PPAR-transcriptionally controlled genes. Future studies should measure the response in the presence of PPARα and PPARγ agonists and antagonists to provide further evidence of this mechanism of action.

In conclusion, this study found that *in vitro* exposure of antral follicles to BzP, BuP, and MBuTP at doses relevant to human exposure disrupted folliculogenesis and steroidogenesis in the mouse ovary. Many of the impacts on follicle growth, gene expression, and hormone levels were similar to legacy phthalates, although, like phthalates, structures within the same class did not necessarily cause the same changes. The similarities between parabens, terephthalate metabolites, and phthalates in ovarian response warrant further study of all three classes of compounds as endocrine disruptors. Until predictive toxicity and substitution selection involve endocrine disruption-sensitive endpoints, introducing replacements with similar structures will have high likelihood of similar endocrine disrupting properties. Given the ubiquitous nature of parabens in global consumer markets, further investigation into the impacts of low dose exposure is essential to inform regulatory standards and safeguard ovarian health.

## Abbreviations

BzP: Benzyl paraben
BuP: Butyl paraben
EDC: Endocrine disrupting chemical
PPAR: Peroxisome proliferator-activated receptor
MBzP: Mono benzyl phthalate
MBzTP: Mono benzyl terepthalate
MuBP: Mono butyl phthalate
MBuTP: Mono butyl terepthalate
DMSO: Dimethyl sulfoxide
ELISA: Enzyme linked immunosorbent assay

## 7. Declaration of generative AI and AI-assisted technologies in the manuscript preparation process

Generative AI was not used in the preparation of this manuscript.

## 8. Author contributions statement

**Courtney Potts**: investigation, project administration, validation, visualization, writing - original draft, writing - reviewing and editing; **Arena Chowdhury**: investigation, visualization, writing - original draft, writing - reviewing and editing; **Raulle Reynolds**: investigation, validation, writing - original draft, writing - reviewing and editing; **Joao Agudo**: investigation, validation, visualization, writing – original draft; **Stephanie Martinez**: investigation; **Hanin Alahmadi**: investigation, **Genoa Warner**: conceptualization, methodology, validation, formal analysis, investigation, writing - original draft, writing - review and editing, visualization, supervision, project administration, funding acquisition

