## Supplemental Information for "Parabens Impair Mouse Antral Follicle Growth and Alter Ovarian Hormone and Gene Expression Similar to Phthalates"

**Contents:**

**Table S1**: DNA oligonucleotide primers used for qPCR gene expression assays

**Figure S1**: Effects of 96 hr *in vitro* BzP (A) and BuP (B) treatment on apoptosis regulator gene expression in antral follicles.

**Figure S2:** Effects of 96 hr *in* *vitro* MBuTP treatment on apoptosis regulator gene expression in antral follicles.

**Figure S3:** Effects of 96 hr *in vitro* MBuTP treatment on steroidogenic enzyme gene expression (A) and sex steroid hormone levels (B) in antral follicles.

| **Table S1**: DNA oligonucleotide primers used for qPCR gene expression assays | | |
| --- | --- | --- |
| **Gene** | **Gene Name** | **Primer Sequence** |
| *Angptl4* | Angiopoietin-like 4 | Forward: 5'-CATCCTGGGACGAGATGAACT-3' |
|  |  | Reverse: 3'-TGACAAGCGTTACCACAGGC-5' |
| *Bad* | B cell leukemia/lymphoma 2 | Forward: 5'-TGAGCCGAGTGAGCAGGAA-3' |
|  |  | Reverse: 3'-GCCTCCATGATGACTGTTGGT-5' |
| *Bax* | Bcl1-associated X protein | Forward: 5'-AGACAGGGGCCTTTTTGCTAC-3' |
|  |  | Reverse: 3'-AATTCGCCGGAGACACTCG-5' |
| *Bcl2* | B cell leukemia/lymphome 2 | Forward: 5'-GCTACCGTCGTGACTTCGC-3' |
|  |  | Reverse: 3'-CCCCACCGAACTCAAAGAAGG-5' |
| *Bcl2l10* | BCL2 like 10 | Forward: 5'-CCACTGCATGAACGCACTAGA-3' |
|  |  | Reverse: 3'-GAGAGCAACTTATCTGCCATCTG-5' |
| *Bid* | BH3 interacting-domain death agonist | Forward: 5'-GCCGAGCACATCACAGACC-3' |
|  |  | Reverse: 3'-TGGCAATGTTGTGGATGATTTCT-5' |
| *Casp3* | Caspase 3 | Forward: 5'-CTCGCTCTGGTACGGATGTG-3' |
|  |  | Reverse: 3'-TCCCATAAATGACCCCTTCATCA-5' |
| *Casp8* | Caspase 8 | Forward: 5'-TGCTTGGACTACATCCCACAC-3' |
|  |  | Reverse: 3'-GTTGCAGTCTAGGAAGTTGACC-5' |
| *Ccna2* | Cyclin A2 | Forward: 5'-AAGAGAATGTCAACCCCGAAAAA-3' |
|  |  | Reverse: 3'-ACCCGTCGAGTCTTGAGCTT-5' |
| *Ccnd2* | Cyclin D2 | Forward: 5'-GAGTGGGAACTGGTAGTGTTG-3' |
|  |  | Reverse: 3'-CGCACAGAGCGATGAAGGT-5' |
| *Cdk2* | Cyclin-dependent kinase 2 | Forward: 5'-ATGGAGAACTTCCAAAAGGTGG-3' |
|  |  | Reverse: 3'-CAGTCTCAGTGTCGAGCCG-5' |
| *Cdk4* | Cyclin-Dependent Kinase 4 | Forward: 5'-ATGGCTGCCACTCGATATGAA-3' |
|  |  | Reverse: 3'-TGCTCCTCCATTAGGAACTCTC-5' |
| *Cdkn1a* | Cyclin-dependent Kinase Inhibitor 1a | Forward: 5'-CCTGGTGATGTCCGACCTG-3' |
|  |  | Reverse: 3'-CCATGAGCGCATCGCAATC-5' |
| *Cdkn2a* | Cyclin-dependent kinase inhibitor 2a | Forward: 5'-CGCAGGTTCTTGGTCACTGT-3' |
|  |  | Reverse: 3'-TGTTCACGAAAGCCAGAGCG-5' |
| *Cdkn2b* | Cyclin-dependent kinase 4 inhibitor B | Forward: 5'-CCCTGCCACCCTTACCAGA-3' |
|  |  | Reverse: 3'-GCAGATACCTCGCAATGTCAC-5' |
| *Cdkn2d* | Cyclin-dependent kinase 4 inhibitor D | Forward: 5'-GGAGCTGGTGCATCCTGACGC-3' |
|  |  | Reverse: 3'-TGGCACCTTGCTTCAGGAGCTC-5' |
| *Cd36* | CD36 molecule | Forward: 5'-ATGGGCTGTGATCGGAACTG-3' |
|  |  | Reverse: 3'-TTTGCCACGTCATCTGGGTTT-5' |
| *Cyp11a1* | Cytochrome-P450 cholesterol side-chain cleavage | Forward: 5'-AGGTCCTTCAATGAGATCCCTT-3' |
|  |  | Reverse: 3'-TCCCTGTAAATGGGGCCATAC-5' |
| *Cyp17a1* | Cytochrome P450 Family 17 Subfamily A member | Forward: 5'-AGTGCTCGTGAAGAAGGGGA-3' |
|  |  | Reverse: 3'-TTTCCTTGGTCCGACAAGAGG-5' |
| *Cyp19a1* | Cytochrome P450 Family 19 Subfamily A member 1 | Forward: 5'-AACCCCATGCAGTATAATGTCAC-3' |
|  |  | Reverse: 3'-AGGACCTGGTATTGAAGACGAG-5' |
| *Fabp4* | Fatty acid binding protein 4, adipocyte | Forward: 5'-TGAAATCACCGCAGACGACAGG-3' |
|  |  | Reverse: 3'-GCTTGTCACCATCTCGTTTTCTC-5' |
| *Hsd17b1* | Hydroxysteroid 17-beta dehydrogenase 1 | Forward: 5'-AAGCGGTTCGTGGAGAAGTAG-3' |
|  |  | Reverse: 3'-ACTGTGCCAGCAAGTTTGCG-5' |
| *Hsd3b1* | Hydroxy-delta-5-steroid dehydrogenase, 3 beta- and steroid delta-isomerase 1 | Forward: 5'-CAGGAGAAAGAACTGCAGGAGGTC-3' |
|  |  | Reverse: 3'-GCACACTTGCTTGAACACAGGC-5' |
| *Ki67* | Antigen identified by monoclonal antibody Ki 67 | Forward: 5'-ATCATTGACCGCTCCTTTAGGT-3' |
|  |  | Reverse: 3'-GCTCGCCTTGATGGTTCCT-5' |
| *Plin1* | Perilipin 1 | Forward: 5'-CTGTGTGCAATGCCTATGAGA-3' |
|  |  | Reverse: 3'-CTGGAGGGTATTGAAGAGCCG-5' |
| *Bactin* | Beta-actin | Forward: 5'-GGGCACAGTGTGGGTGAC-3' |
|  |  | Reverse: 3'-CTGGCACCACACCTTCTAC-5' |
| *Star* | Steroidogenic Acute Regulatory | Forward: 5'-CAGGGAGAGGTGGCTATGCA-3' |
|  |  | Reverse: 3'-CCGTGTCTTTTCCAATCCTCTG-5' |

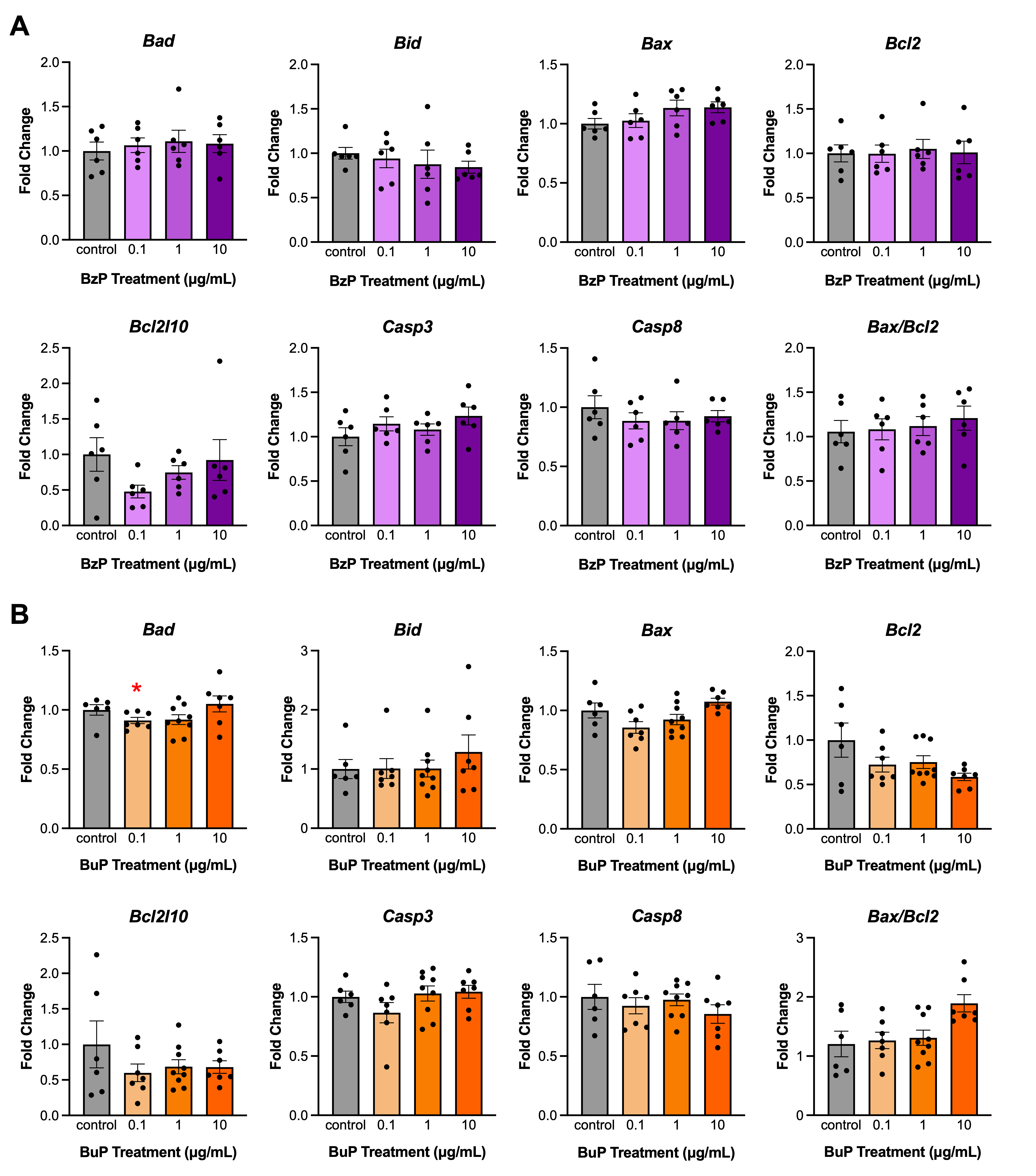

**Figure S1**: Effects of 96 hr *in vitro* BzP (A) and BuP (B) treatment on apoptosis regulator gene expression in antral follicles. All gene expression is relative to the housekeeper gene *B-actin*. Graphs represent means ± SEM from 6-7 experiments per treatment group. Asterisks (∗) indicate significant differences from the control (p ≤ 0.05).

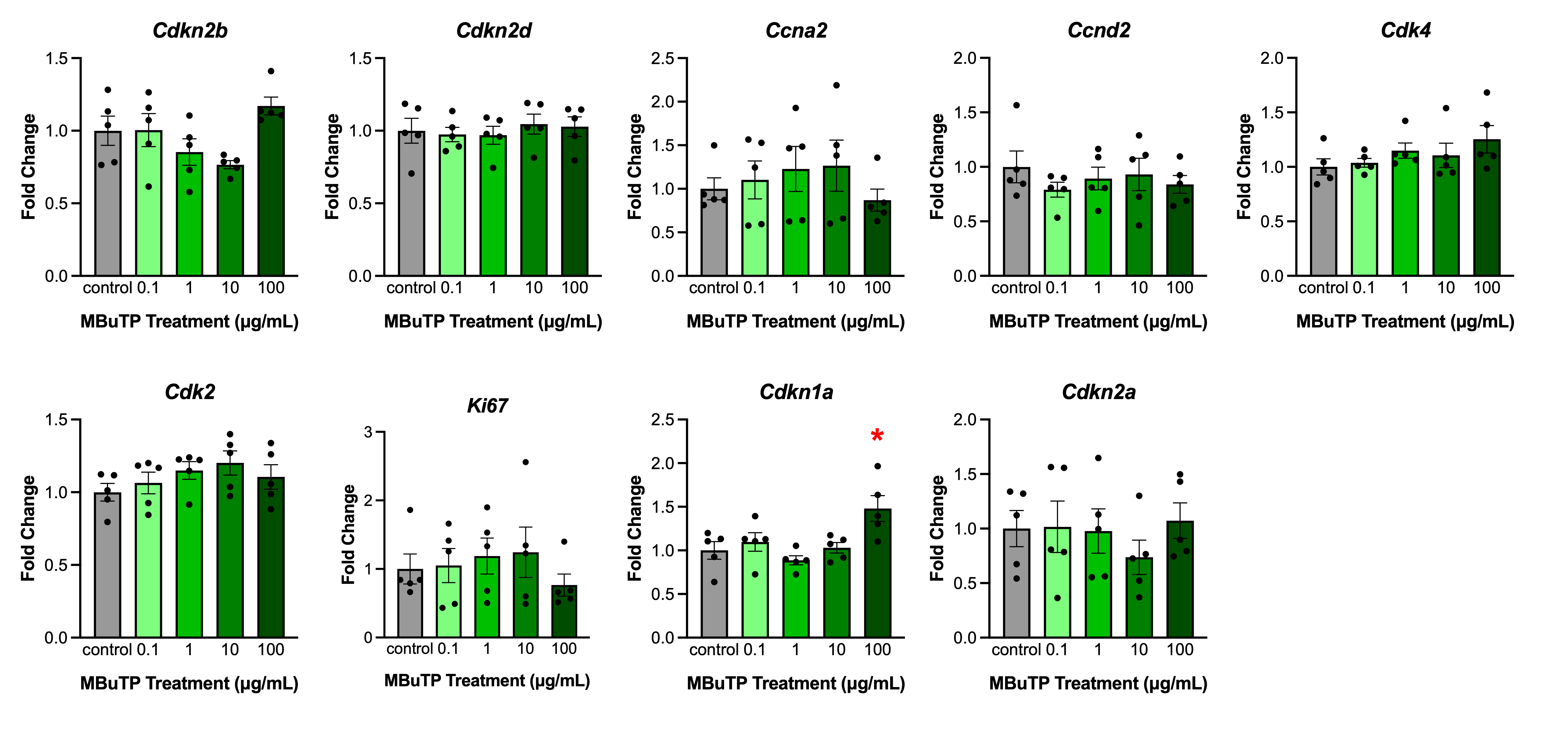

**Figure S2:** Effects of 96 hr *in* *vitro* MBuTP treatment on apoptosis regulator gene expression in antral follicles. Graphs represent means ± SEM from 5 experiments per treatment group. Asterisks (∗) indicate significant differences from the control (p ≤ 0.05).

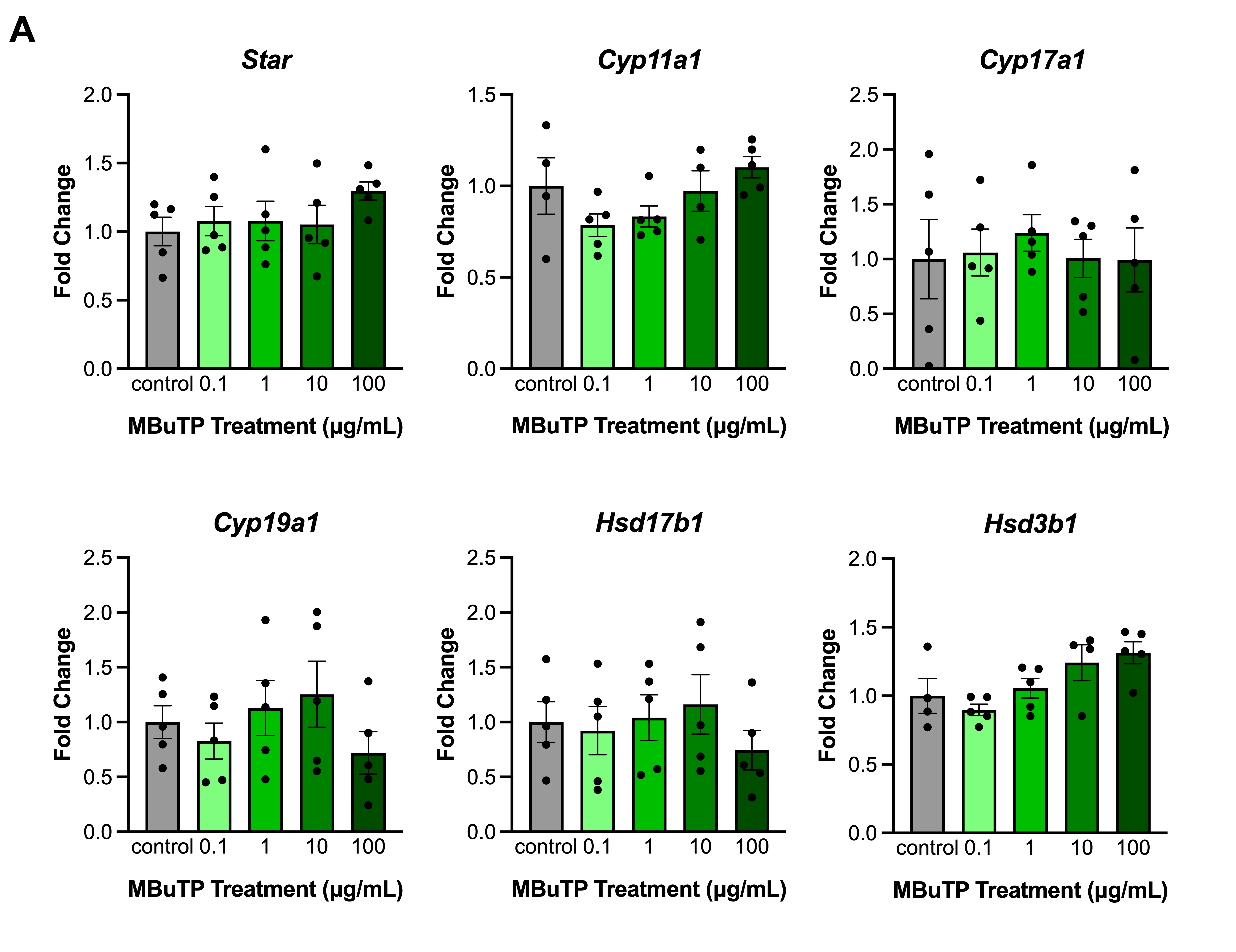

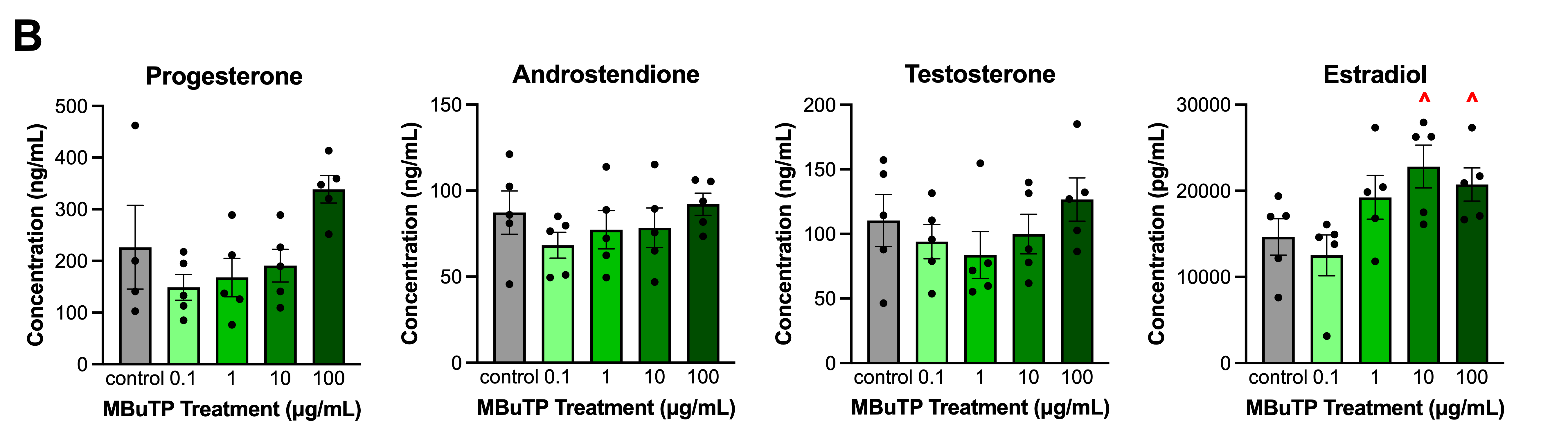
**Figure S3:** Effects of 96 hr *in vitro* MBuTP treatment on steroidogenic enzyme gene expression (A) and sex steroid hormone levels (B) in antral follicles. All gene expression is relative to the housekeeper gene *B-actin*. Graphs represent means ± SEM from 5 experiments per treatment group. Asterisks (∗) indicate significant differences from the control (p ≤ 0.05) and carets (^) indicates a trend toward significance (p ≤ 0.10).
